# Differential rapid plasticity in auditory and visual responses in the primarily multisensory orbitofrontal cortex

**DOI:** 10.1101/869446

**Authors:** Sudha Sharma, Sharba Bandyopadhyay

## Abstract

In a dynamic environment with rapidly changing contingencies, the orbitofrontal cortex (OFC) guides flexible behavior through coding of stimulus value. Although stimulus-evoked responses in the OFC are known to convey outcome, baseline sensory response properties in the mouse OFC are poorly understood. To understand mechanisms involved in stimulus value/outcome encoding it is important to know the response properties of single neurons in the mouse OFC, purely from a sensory perspective. Ruling out effects of behavioral state, memory and others, we studied the anesthetized mouse OFC responses to auditory, visual and audiovisual/multisensory stimuli, multisensory associations and sensory-driven input organization to the OFC. Almost all, OFC single neurons were found to be multisensory in nature, with sublinear to supralinear integration of the component unisensory stimuli. With a novel multisensory oddball stimulus set, we show that the OFC receives both unisensory as well as multisensory inputs, further corroborated by retrograde tracers showing labeling in secondary auditory and visual cortices, which we find to also have similar multisensory integration and responses. With long audiovisual pairing/association, we show rapid plasticity in OFC single neurons, with a strong visual bias, leading to a strong depression of auditory responses and effective enhancement of visual responses. Such rapid multisensory association driven plasticity is absent in the auditory and visual cortices, suggesting its emergence in the OFC. Based on the above results we propose a hypothetical local circuit model in the OFC that integrates auditory and visual information which participates in computing stimulus value in dynamic multisensory environments.

**Significance Statement:** Properties and modification of sensory responses of neurons in the orbitofrontal cortex (OFC) involved in flexible behavior through stimulus value/outcome encoding are poorly understood. Such responses are critical in providing the framework for the encoding of stimulus value based on behavioral context while also directing plastic changes in sensory regions. The mouse OFC is found to be primarily multisensory with varied nonlinear interactions, explained by unisensory and multisensory inputs. Audio-visual associations cause rapid plastic changes in the OFC, which bias visual responses while suppressing auditory responses. Similar plasticity was absent in the sensory cortex. Thus the observed intrinsic visual bias in the OFC weighs visual stimuli more than associated auditory stimuli in value encoding in a dynamic multisensory environment.

## Introduction

Function of the OFC, a part of the prefrontal cortex (PFC), encompasses the broad representation or model of an animal’s sensory environment and relevant actions and their relationship to outcomes (Wilson et al., 2014). A large fraction of OFC neurons encodes the sensory attributes and subjective value of outcomes associated with external stimuli (Padoa-Schioppa and Conen, 2017). Stimulus-evoked signals in the OFC can convey the relative value of significant outcomes, distinguishing between appetitive and aversive outcomes (Morrison and Daniel Salzman, 2011). The neurophysiological properties of the OFC along with its connections with the sensory and limbic structures place the OFC in a unique position (Carmichael and Price, 1995), where it can potentially use different sensory inputs to integrate and modulate them to perform value computation and constant updating based on dynamic needs. Although studies of the function of the OFC have mainly used unisensory stimuli (Sadacca et al., 2018), the real sensory world stimuli are mostly combinations of multiple modalities, either modulating each other or are associated with each other. Many recent behavioral studies clearly indicate the importance of multisensory integration and modulation in behavior (Raposo et al., 2012)(Raposo et al., 2014). Thus to understand the function of the OFC in assigning value to stimuli to achieve behavioral flexibility, it is important to consider the circuits involved in providing multisensory information to the OFC, representation of multisensory stimuli in the OFC and how the representations may change with associations, purely from a sensory point of view. Such understanding provides the crucial background representation of the sensory world on top of which, reward-punishment (O’Doherty et al., 2001), action-outcome (Simon et al., 2015) and prediction error (Schultz, 2016) signals operate to achieve goal-directed behavior in a seamless manner. Further, the outputs of the OFC based on such multisensory information can potentially also modulate representation in lower order primary sensory regions (Winkowski et al., 2013) to aid behavior.

Most multisensory representation studies, primarily audiovisual, of the OFC and PFC, are based on nonhuman primates, showing likely domain specificity (Romanski, 2004) in the frontal cortex. However, later studies and more recent studies favor a category free population or mixed selectivity of neurons in the PFC. Higher cortical areas have neurons that are deployed in different ways for specific task features and evolve according to task demands(Raposo et al., 2014). However, a detailed anatomical and physiological study of the mouse OFC in encoding different sensory inputs and integrating them in its unique way to extract the best possible estimate of the environment remains to be undertaken.

In this paper, using auditory and visual stimuli, we study the underlying multisensory representation and multisensory response properties in the mouse OFC. We show that auditory, visual and audiovisual (henceforth referred to as multisensory) stimuli have different representations and response properties within the same neurons of the OFC, with almost all neurons studied being multisensory in nature. With retrograde labeling studies, we show the possible origins (direct and indirect) of auditory and visual inputs to the OFC. The origins of multisensory inputs lie in nonprimary auditory and visual cortices which we show have multisensory responses, with similar multisensory modulation of responses as observed in the OFC. Using a novel multisensory oddball paradigm we also show that the OFC receives exclusive unisensory (auditory and visual) as well as exclusively multisensory inputs. Finally, we show that long auditory-visual associations induce rapid plasticity in the OFC, which is differential in nature, showing strong suppression to auditory responses with an effective enhancement of visual responses. Such association driven rapid plasticity is absent in the sensory cortices, which provide input to the OFC and could thus be emergent in the OFC. The observed visual bias in the OFC due to multisensory associations has a number of implications in understanding goal-directed flexible behavior in a realistic multisensory environment and in understanding neuropsychiatric disorders associated with OFC dysfunction.

## Materials and Methods

All procedures were approved by the Indian Institute of Technology, Institutional Animal Care and Use Committee. Animals were maintained in 12 hr light 12 hr dark cycle and all experiments were performed during the dark cycle on the C57 strain from Jackson laboratories.

### Anatomy

Mice aged more than postnatal day 60 (> P60, male or female) were anesthetized with isoflurane (5% induction and 1.5% maintenance) and placed on a stereotaxic frame. The body temperature was kept at 37 °C throughout the procedure using a heating pad. An incision was made to expose the skull. A burr hole (~0.5 mm diameter) was made above the injection site. Tracers were loaded in a glass micropipette mounted on a Nanoject II attached to a micromanipulator and then injected at a speed of 20 nL per minute. Retrobead injections were targeted stereotactically into the left OFC using the following co-ordinates(Paxinos and Franklin, n.d.): +2.5 mm from the bregma point and 1 from the midline at a depth of 1.8mm. After recovery from anesthesia, mice were left in their home cages for 2 weeks following the injection.

#### Histology and quantification

Mice were deeply anesthetized with isoflurane, perfused transcardially with 0.1M phosphate-buffered saline followed by 4% paraformaldehyde. Brains were harvested and after a post-fixation period of 8-10 hours in 4 °C, 100 um thick sections were cut using a vibratome (Leica VT1000S) and images were taken in a fluorescence microscope (Leica DM2500). Brain slices encompassing the span of auditory cortex (ACX) rostro-caudally were selected, areas were demarcated based on a mouse brain atlas(Paxinos and Franklin, n.d.) and labeled cell bodies were counted manually in each of the different regions on all sections.

### Electrophysiological recordings

Mice (58 mice P31 to P40 and 2 mice P27 and P28) were anesthetized with isoflurane (5% induction and 1% maintenance) and the skull was attached to a stainless steel plate for fixing the head of the animal for inserting the electrodes and performing recordings. Animals were kept warm throughout the experiment by maintaining an external body temperature of 37°C throughout the procedure using a heating pad. An incision was made to expose the skull. A craniotomy (~2 mm diameter) was made above the left OFC (or ACX) for recordings (+2.5mm from bregma and +1 mm from the midline, co-ordinates of ACX, (Winkowski et al., 2013). Extracellular recordings were performed using tungsten Microelectrodes Array (MEA) of impedance 3-5 Megaohms (MicroProbes, USA). 4×4 custom-designed metal MEAs with an inter-electrode spacing of 125 microns were used. The array was advanced slowly to a depth of 1800 um from the surface (Fig. 1A) into the OFC using a micromanipulator (MP-285, Sutter Instrument Company, Novato, CA). The electrodes were allowed to settle for >20 min before the stimulus presentation was started. Signals were acquired after passing through a unity gain head stage (Plexon, HST16o25) followed by PBX3 (Plexon) preamp with gain of X1000, to obtain the wideband signal (used to extract LFP, 0.7 Hz to 6 kHz) and spike signals (150 Hz to 8 kHz) in parallel and acquired through National Instruments Data Acquisition Card (NI-PCI-6259) at 20 kHz sampling rate, controlled through custom-written MATLAB (Mathworks, Natick, USA) routines. Further, all online/offline analysis was performed using custom-written MATLAB routines. After the experiment was completed the brain of the animal was routinely isolated and kept in a 4% solution of PFA for later visualization of the site of recording in 100 um thick sections through cresyl violet staining.

**Figure 1.**
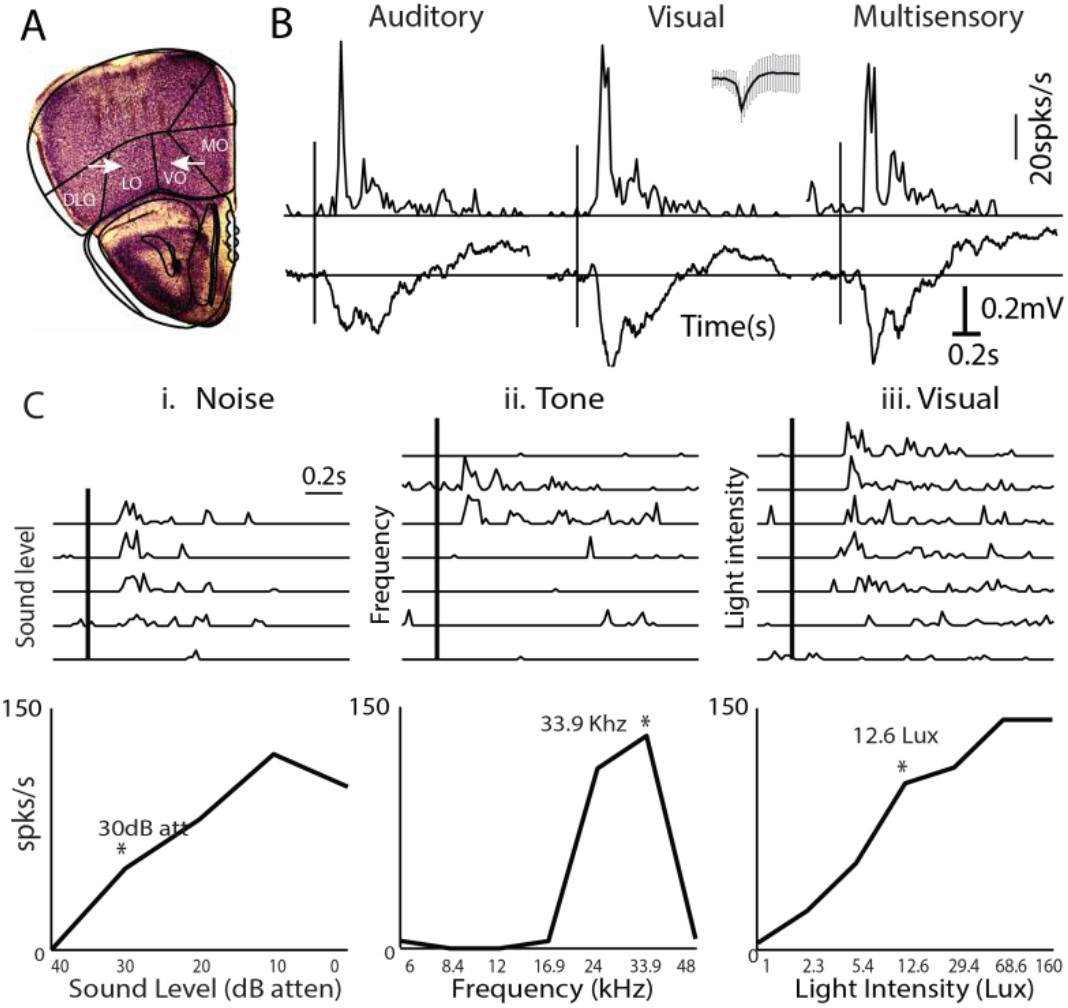
Multisensory responses of mouse OFC single neurons. **A.** Left: Example of electrode tracks (arrowheads) shown in Nissl stains of brain slices showing recording locations in LO/VO. **B**. Representative PSTHs of a single OFC neuron in response to auditory (tone= 33.9 kHz), visual (LED blink 12.6 lux) and multisensory stimuli. *Inset*: spike shape. Mean LFPs on the same channel are shown below each PSTH. **Ci-iii.** (Top row) PSTHs of single neurons in response to multiple sound levels of a noise burst (i), multiple frequencies (ii) and multiple intensities of LED blink (iii) are shown. (Bottom row) The mean rate response as a function of sound level, frequency and light intensity are plotted. Asterisks represent the chosen values presented for the *M* stimulus in each case.

### Auditory and Visual Stimulation

Sound stimuli were presented to the ear contralateral to the recording hemisphere inside a soundproof chamber (IAC Acoustics, IL, USA), 10 cm away from the right ear (contralateral) of the mouse, with TDT electrostatic speakers (ES1) driven by TDT drivers ED1 after attenuation by TDT attenuators (PA5), generated through TDT RX6 using custom software written in MATLAB. The acoustic calibrations, performed with microphone 4939 (Brüel&Kjær, Denmark), of the ES1 speakers (TDT) in the sound chamber, showed a typical flat (+/− 7 dB) calibration curve from 4-60 kHz. 0-dB attenuation on PA5 corresponded to a mean (across frequency) of 95 dB SPL. Sound stimuli consisted of pure tone (6-48 kHz) or white noise (bandwidth 6-48 kHz) bursts. After usual initial characterization of single units with different intensity noise bursts (5 repetitions, 50ms duration, 5ms rise and fall), single-unit responses to tones of different frequencies (5 repetitions each, 6-48 kHz, ½ octaves apart, 50ms duration with 5ms rise and fall, 5s inter stimulus interval at ~65-75 dB SPL, usually ~10-20 dB louder than the usual noise threshold) were collected. A particular frequency was decided based on the maximum number of responsive channels to the different tones for further experiments and for audiovisual stimulation (stimulus *A*). Single unit spike times were obtained from the acquired spike channel data using threshold crossing and spike sorting with custom-written software in MATLAB. Responses to *A* with 30 repetitions were collected following the characterization and used for further analysis.

For visual stimulation, a white LED (5 mm Round White LED(T-1 3/4)) kept 5 cm away from the eye of the animal was used. Full-field illumination to the contralateral eye was provided and the intensity was varied through the NI DAQ card, using MATLAB routines. LED blink stimulus was presented for 10 ms (*V* stimulus) with an interstimulus interval of 5s in order for the cones in the retina to recover. LED flash of 10 ms duration allowed for response rates of the *A* and *V* to be comparable. A petroleum jelly based eye ointment was applied in order to prevent the eye from drying. Single unit rate responses were obtained for varying intensities of the LED (1-160 lux, 5-7 steps, Model: LX1010b, Digital Lux Meter, 0 to 20000 lux) and threshold intensity was determined. For further experiments and audiovisual stimulation, a chosen intensity above the threshold (of stimulus *V*) was used, which corresponded to 5-30 lux. As with the *A* stimulus, responses to *V* with 30 repetitions were collected.

Bimodal stimulation and audio-visual pairing comprised of simultaneous presentation of the *A* and *V* stimuli (*M*), as above. 100 repetitions of the *M* stimuli were presented and the first 30 were considered for calculating the *M* response for comparison with responses to unisensory stimuli. Following the 100 presentations of the *M* stimulus, considered as the *A* and *V* pairing, 30 repetitions of *A* and then *V* stimuli were presented to obtain the effect of audiovisual pairing. For comparisons between before and after pairing, the 2 mice aged less than 31 days were not included due to possible immaturity in the visual system. Further, for the pairing data set only units with the same spike shapes present in all 5 sets of responses (A-before, V-before, M, A-after and V-after) were included. Thus, of the total 158 units obtained in 12 animals with responses to *A*, *V* and *M*, 91 units were used for the analysis of the effect of pairing.

### Data Analysis

Spike Sorting was done offline in custom-written MATLAB scripts. Data were baseline corrected and notch filtered (Butterworth 4^th^ order) to reject any remnant power supply 50Hz oscillations. Wideband data were bandpass filtered for local field potentials between 1 and 500 Hz and spiking activity was obtained directly from the spike channels of the PBX3 preamp. Deviations above 4 standard deviations from the baseline were isolated and based on shapes, spike waveforms were clustered into different groups.

#### Calculation of latencies, spike rate, PSTH and response duration

For data from OFC, a moving window of size 100 ms in steps of 20 ms (ACX, 50 ms window in steps of 5 ms) after the stimulus start was used for comparison with a random 100 ms window from the baseline (between 400 ms to 10 ms preceding stimulus onset) for detecting a significant response (*p*<0.05, *paired t-test*). The middle of the first significant window was taken as the latency of response. The time bin (20 ms bins for OFC, 5 ms for ACx) corresponding to the maximum spike rate within 500 ms of stimulus onset was detected. Mean spike rate in a 100 ms (ACx: 25ms for A and V, 50ms for M) window around the peak was taken as the spike rate. All peristimulus time histograms (PSTHs) shown were calculated based on 20 ms bins (5 ms bins for ACx) by averaging the firing rate from multiple repetitions of the same stimulus. A sliding response window of size 100 ms (50 ms for ACX), starting from the stimulus start, in 20 ms steps (5 ms for ACX), was compared with the random 100 ms window in the baseline for significance. Consecutive significant bins with a time difference of less than 100 ms (25 ms for ACX) between were joined together in the response for the determination of response duration. The LFP signal was converted to root mean square (*rms*) in the same time windows used for spike data and significance was calculated in a similar manner as above. The same procedure was followed for both OFC and ACX data as described above for the case of spikes. The LFP response strength was calculated as the *rms* of the average LFP waveform in a 100 ms window centered around the maximum dip in LFP following stimulus onset.

#### Classification of neurons

Neurons that responded both to auditory and visual stimulation or showed significant change on presentation of multisensory stimulus when compared to the larger of the unisensory responses are considered as multisensory units. Neurons that were responsive to only auditory or visual and do not show any significant change on multisensory stimulus presentation are considered as exclusively auditory or exclusively visual units respectively. To calculate percent change on multisensory presentation, Multisensory Enhancement Index, *MEI*=[*r*(*M*) − *max*(*r*(*A*)*,r*(*V*)]/*max*(*r*(*A*)*, r*(*V*)) is used, where *r*(*X*) is considered as the rate response to stimulus *X* = *A*, *V* or *M*. For comparison of spike rates to the auditory and visual stimulus, Unisensory Imbalance (*UI*) is calculated as (*r*(*V*)−*r*(A))/ (*r*(*V*)+*r*(A)) (Miller et al., 2017).

#### Comparisons of slopes

First a straight line through the origin (*y*=*mx*) was fit using minimum mean squared error, to the scatter data of after versus before responses and the slope was estimated. For comparisons of slopes of the best fit lines to scatter plots of after pairing responses versus before pairing responses across conditions, bootstrap analysis(Efron et al., n.d.) was performed to obtain confidence intervals on the slopes. Bootstrap populations (*n*=1000) were created by randomly sampling before-after pairs of responses with replacement from the original dataset. We obtained 1000 slope values from the population of 1000 bootstrap sets. The slopes from two different conditions were considered to be significantly different if the 95% confidence intervals of the two did not overlap with each other and accordingly for one-sided comparisons.

## Results

### Single neurons in the mouse OFC are primarily multisensory

In order to investigate the relative prevalence of multisensory neurons and interaction of two unisensory stimuli in the mouse OFC, we first characterize responses of single neurons to an auditory, a visual and a multisensory stimulus (Fig. 1B). Single unit recordings (Methods) were performed in the anesthetized mouse OFC. Recording site depth and the location were determined post hoc by cresyl violet staining of the coronal sections of the mouse brain (Figure 1A). Usually, intensity of Presentation above noise threshold was first determined based on responses to broadband noise at different intensities (Fig. 1Ci, asterisk) and then tuning of single units was obtained with tone presentations (Fig. 1Cii, Methods) at 65-75 dB SPL. A particular frequency (Fig. 1Cii, asterisk) was chosen for the auditory component (*A*) of the multisensory stimulus (*M*). The intensity of the visual component (*V*) of the subsequent multisensory stimulus was determined based on responses of single units to the visual stimulus at multiple intensities (Fig. 1Ciii, asterisk, Methods). Overall, data from 158 single units in the mouse OFC across 12 animals were analyzed for unisensory and multisensory response characterization. These units showed a significant response to at least one of the three stimuli, *A*, *V* and *M*.

Most neurons (96.2%, n=152/158) in the OFC were found to be multisensory in nature; 147 units responding to auditory as well as visual stimuli and 5 units did not respond to both *A* and *V* but were multisensory in nature (2 responded to only *A*, and 2 to only *V* and suppressed to *M* and 1 unit responded to neither *A* nor *V* but responded to *M*). Representative response PSTHs (20 ms bin) to each of the three stimuli *A*, *V*, and *M*, for a multisensory neuron, are shown in Fig. 1B. Local field potentials (LFPs) recorded simultaneously from the same electrode are shown below the PSTHs, which also show similar characteristics. The remaining 6 units were found to be exclusively auditory (same response to *M* as to *A*) and none were exclusively visual. Similar to early sensory cortices, neurons in the OFC were also frequency selective (Figure 1Cii), and these neurons were found to be sensitive to increases in the intensity of the auditory and visual stimuli (Fig. 1Ci&iii). Intensities of *A* and *V* chosen for the presentation of *M* were within the general dynamic range of the neurons and not at saturating activity levels.

We further characterize the multisensory responses of OFC single units based on the modulation of unisensory responses because of the simultaneous presence of a stimulus of another modality, in the multisensory stimulation. Almost equal proportions of neurons showed significant modulation by *M* (45.3% 69/152, Fig. 2) and no modulation by *M* (54.3%, 83/152). The proportions of the two types of neurons were not significantly different (test of proportions). The neurons with no modulation by *M* responded to both *A* and *V* but their rate responses to *M* were not significantly different from the larger of the responses to the unisensory stimuli (example neuron, Fig. 2Ai). Thus, although the response rates to each of the unisensory stimuli (*A* and *V*) were individually not saturating, presentation of the two stimuli together did not significantly alter the response strengths. Thus these neurons effectively show nonlinear interactions between the two stimuli when presented simultaneously. Out of the remaining 45.3% neurons that significantly altered their spike rates on presentation of *M* compared to the unisensory conditions, a significantly larger proportion of neurons (41 units, 26.9%, *p*<0.01, binomial test) showed suppression of responses (example neuron, Fig. 2Bi) compared to enhancement (28 units, 18.3%, example neuron Fig. 2Ci). The corresponding LFPs (shown below each PSTH) collected on the same channels as the units show similar character in terms of enhancement and suppression (bottom row, Fig. 2BC). Population PSTHs of each set of neurons, no modulation (*M0*), suppression (*MS*) and enhancement (*ME*) are shown in Fig. 2A-C, along with population mean LFPs, in response to the 3 stimuli (columns ii-iv) below the population PSTHs. To quantify the degree of modulation, the Multisensory Enhancement Index (*MEI*, Methods) was calculated. The scatter plot of *MEI* of each unit versus the unisensory imbalance (*UI*, Methods) between the rate responses to *A* and *V* (Fig. 2D) show that most suppressed units had a high absolute *UI*, while enhanced units had lower *UI*. The median abs(*UI*) of the 3 classes of multisensory modulation were significantly different (*ranksum test*) from each other (no change *n_ei_*= 0.24, enhanced *e_ei_*= 0.12 and suppressed *s_ei_* = 0.43, Fig. 2E) Thus, unisensory imbalance appears to be a determining factor of whether a neuron in the OFC would be suppressed or enhanced or show no changes. The median abs(*UI*) values for units with larger auditory (*LA*) responses (negative *UI*) was higher (*p*<0.05, *ranksum*) than units with larger visual (*LV*) responses, indicating OFC neurons at the chosen light intensity of *V* and frequency and sound level of *A* had stronger responses to *A* than *V*. However, the distribution of observed abs(*UI*) for *LA* and *LV* units were not different (*kstest*, NS). The above indicates that the suppression of units is due to having larger auditory responses which coincide with higher *UI* in this dataset. Units with high positive *UI* (*LV*) do not get suppressed, while larger fraction (p<0.001, test of proportions) of *LA* units gets suppressed. Thus the primary effect of the presentation of *M* is to inhibit the responses of neurons responding strongly to *A*.

**Figure 2.**
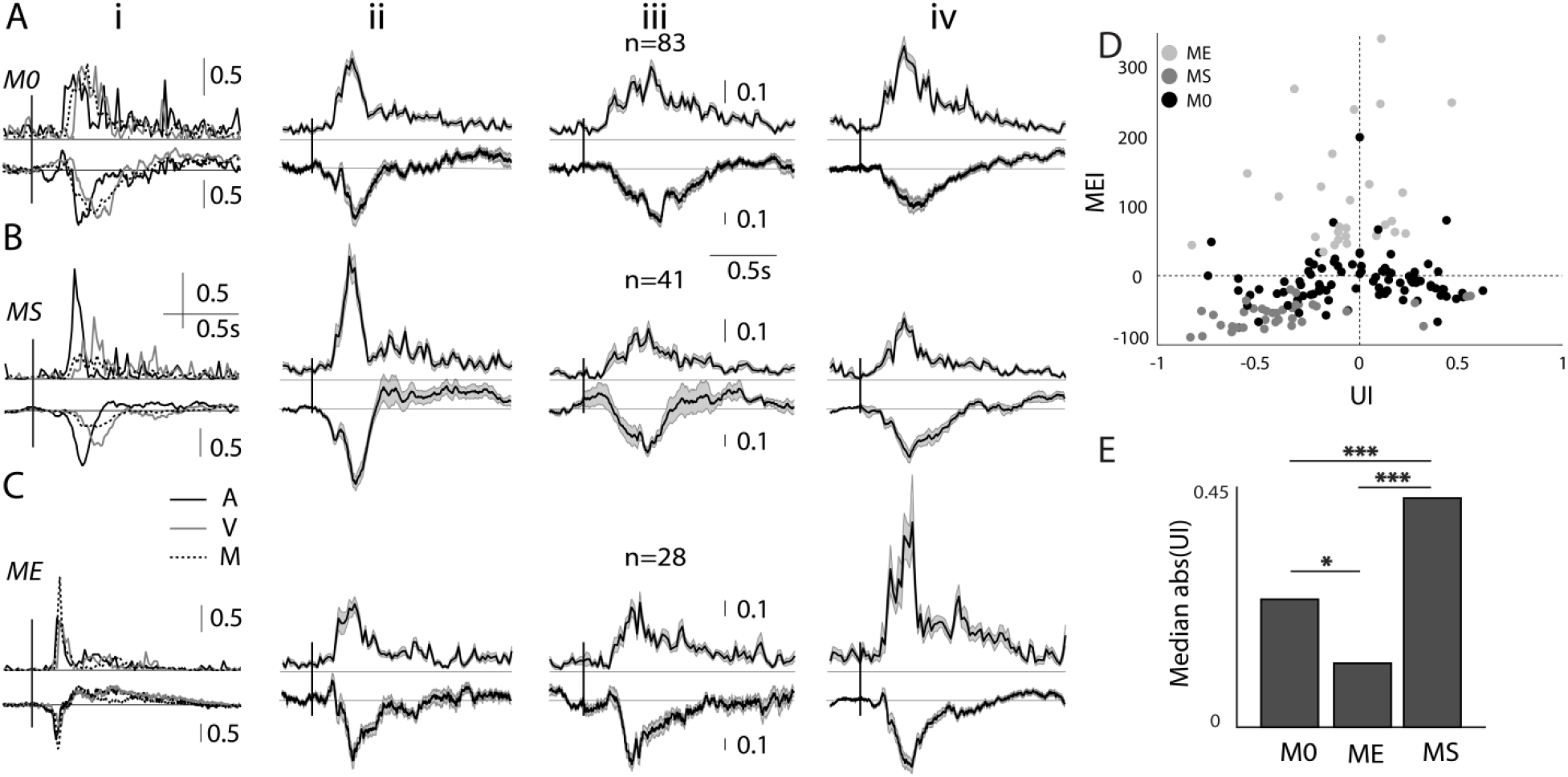
Multisensory modulation of mouse OFC single neurons. A-C. Each row shows representative examples of single unit PSTHs (left column) in response to *A* (black), *V* (gray) and *M* (black dashed) followed by population PSTHs with *SEM* (gray shading)(normalized by the maximum response of the *A* and *V* stimuli) in response to *A* (second column), *V* (third) and *M* (fourth). A: no modulation (*M0*), B: suppressed (*MS*) C: enhanced (*ME*). **D.** Scatter plot of *MEI* versus *UI*, each dot represents a single unit and shade indicates the kind of modulation. **E.** Barplots showing median abs(*UI*) of the groups of 3 kinds of modulation (* p<0.05, *** p<0.001). n represents the number of single neurons, vertical bars over the example neurons and population PSTHs is the length of normalized rate responses (a.u.).

### Response latency and response duration indicate modality-specific coding

As the mean firing rate response of OFC single neurons to each of the unisensory modalities (*A* and *V*) determine multisensory modulation we also investigate how their basic temporal response properties relate to the type of modulation. Figure 3Ai-iii shows the cumulative response distribution functions (CDFs) of response latency of all neurons to the three stimuli *A*, *V* and *M* (i-iii) separated into groups by their modulation index (identified by colors). Comparing the median latency between the 3 stimulation conditions (*A*: 180ms, *V*: 280ms and *M*: 220ms, Fig. 3Aiv) shows that responses to *A* have significantly lower latency than the other two cases, while responses to *V* have significantly higher latency than that of *M*. The response latencies to *V* and *M* were not different in the *ME/MS/M0* types of units. However, *ME* units had significantly longer response latency to *A* than *MS* (p<0.001) and *M0* (p<0.01) units. Thus long latency *A* responses, indicative of weaker auditory drive, were usually enhanced as also concluded from rates above. Thus these observations are further evidence that the strength of auditory drive in OFC units determines the kind of modulation the units have with a multisensory stimulus.

**Figure 3.**
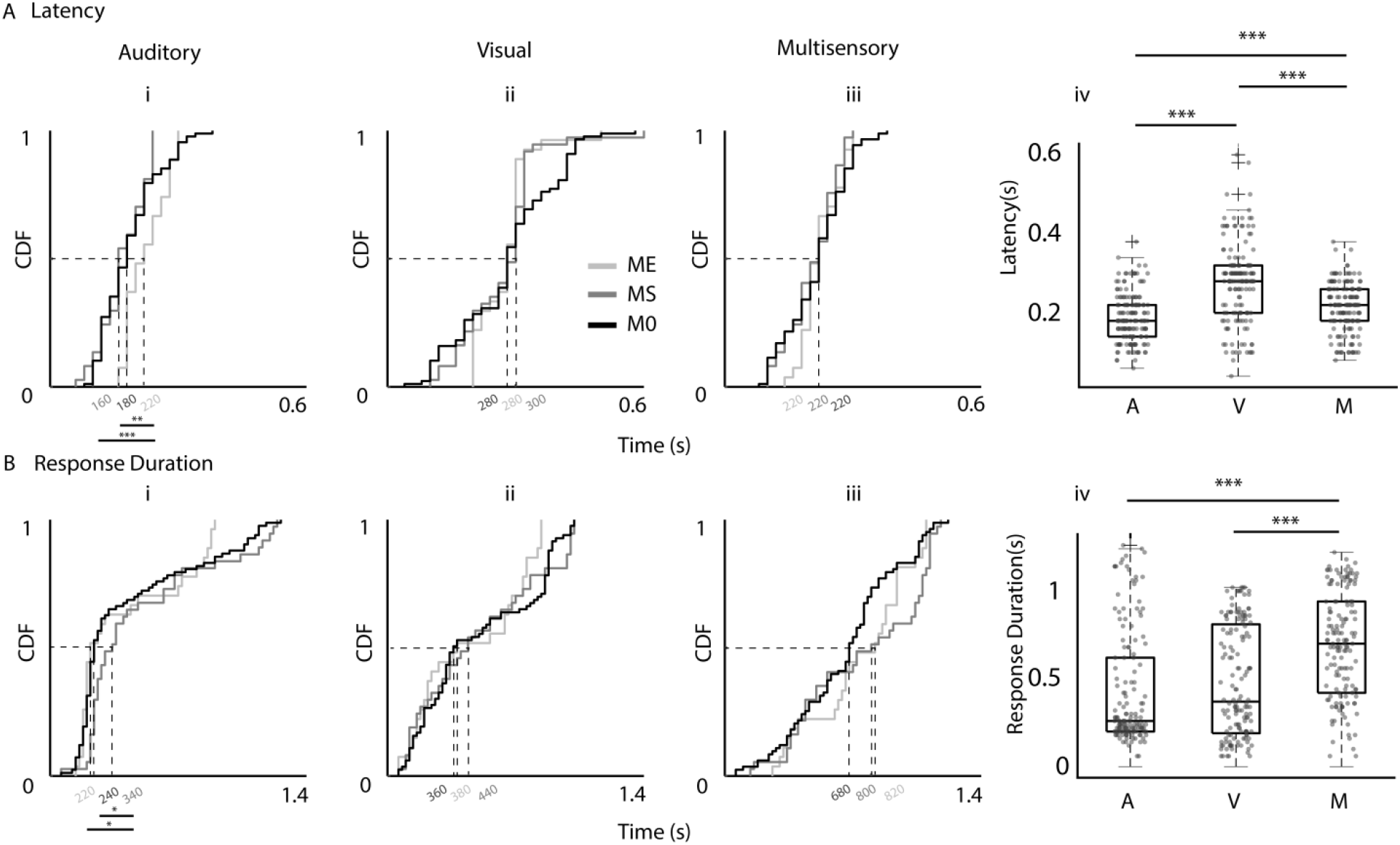
Dependence of latency and response durations of OFC units to *A*, *V*, *M* stimuli. **Ai-iii.** CDF plots of latencies of OFC single units with different types of modulations (identified by shading of lines) to *A*, *V*, and *M* (i-iii) stimuli. **Aiv**. Box plots of latencies of all single units to each of the stimuli, *A, V* and *M*. **Bi-iv.** Same arrangement of CDF plots and box plots as in **A** for response duration.

Unlike in the primary auditory and visual sensory cortices, *A*, *V* and *M* stimuli evoked a considerably long-duration response and LFPs (Fig. 1B&2) in the OFC. Response duration CDFs for each stimulus separated into groups based on their multisensory modulation (Fig. 3Bi-iii) show that in case of the *V* stimulus-response durations were independent of multisensory modulation. However, median *A* response duration was significantly shorter for *ME* (220 ms) units than that of *MS* (340 ms, *p*<0.05), which is significantly longer than that of *M0* (240 ms *p*<0.05). Thus, again, weaker auditory drive, indicated by short duration responses, coincides with multisensory enhancement while stronger auditory drive, indicated by long response duration, coincides with multisensory suppression. The median response duration of OFC single units, to *A*, *V* and *M* stimuli (Fig. 3Biv), show that the *M* stimulus (720ms) evoked significantly longer responses compared to the *A* (260 ms, *p*<0.001) and *V* (380ms, *p*<0.001) stimuli, possibly indicating temporal summation with the simultaneous presentation of two stimuli. It could additionally be also due to release from inhibition from one of the unisensory stimuli (likely auditory, Fig. 2C) within the multisensory response.

Thus based on short response latency and long response duration it is further corroborated that the higher strength of auditory input, as also determined by rates, indicated multisensory suppression. Weak auditory drive indicated by long latency and short duration responses indicated multisensory enhancement. Multisensory modulation is largely independent of the response to the unisensory visual component of the multisensory stimulus.

Our results based on response rates and basic temporal response characteristics suggests stimulus type (*A*, *V*, *M*) specific coding as each of the stimuli have different median latencies of response, different response strengths to *A* and *V*, and longer duration responses to *M* compared to *A* and *V*. Further, simultaneous presentation of *A* and *V* leads to variety of nonlinear interactions between the individual unisensory responses. All the above, especially different response latencies to *A*, *V*, and *M* suggest that each of the three types of stimuli activates partially unique and partially overlapping pathways. To explain the difference in latency between the *A* and *M* stimulus, we hypothesize that neurons in the OFC receive inputs from a separate pathway activated only by multisensory inputs but not by the individual unisensory components, which also likely inhibits the auditory inputs to the OFC. For the above hypothesized model to achieve the latency differences, a low threshold multisensory-activated inhibition on the auditory inputs is required. The suppression by the *M* stimuli in *LA* units provides further evidence in favor of the existence of such an inhibitory input. The same could also be achieved by a pathway that is activated by both an ‘only visual’ and ‘a multisensory’ stimulus. However, the latter is less likely due to the very large difference (~100 ms) in latency between the auditory and visual responses of OFC single units. We test the above based on anatomical and physiological evidence.

### Sources of auditory, visual and multisensory inputs to the mouse OFC

In order to look at the possible sources of auditory, visual and multisensory inputs into OFC, we injected fluorescent retrograde tracers into OFC (see Methods, Fig. 4Ai, white arrow). The cortical region on the rostrocaudal axis encompassing the auditory cortex (Fig. 4Aii), based on anatomical landmarks(Paxinos and Franklin, n.d.), was imaged. Numbers of fluorescently labeled cell bodies (Fig. 4Aiii, *n*=5 animals, see Methods) in different regions along the mediolateral extent up to the rhinal fissure (Fig. 4Aii, arrow) were determined. Regions were demarcated based on the mouse brain atlas(Paxinos and Franklin, n.d.) and included the auditory cortex (ACx): ventral, primary and dorsal, and the visual cortex (VCx): Lateral, Primary and Medial (M/L) divisions. All regions have direct projections to the OFC, with secondary regions showing much stronger labeling than primary regions in both auditory and visual cortices (Fig. 4B). Thus direct anatomical pathways carrying auditory and visual information to the OFC exist (Zingg et al., 2014) and could provide the necessary source of inputs giving rise to both unisensory and multisensory responses in the OFC. Other possibilities also exist, as observed by the retrograde labeling of regions such as TeA, Ect, Prh, PPC, Basolateral nucleus of Amygdala, known to be multisensory(Morrow et al., 2019)(Ghazanfar and Schroeder, 2006)(Lyamzin and Benucci, 2019). Among the auditory and visual sensory cortical regions projecting directly to the OFC, with LFP recordings along the mediolateral axis (Fig. 4C, electrode tracks, white arrows) we find that the dorsal auditory cortex and lateral secondary visual cortex (AuD and V2L) have responses to both auditory and visual (Fig. 4Di, iii) stimuli, with expected latencies (Fig. 4Dii, iii) in primary and secondary regions. Thus the above secondary regions, with their strong projections to the OFC, are the most likely origins of multisensory (A/V) inputs to the OFC. Single unit recordings in the AuD and V2L show multisensory responses in spike rates (Fig. 4E), as observed in the OFC (Fig. 1). Similar fractions of modulatory effects were found in the AuD as in the OFC (86/147, 58.5% *M0*, 23/147, 15.6% *ME* and 38/147, 25.85% *MS*, test of proportion), showing that such modulatory effects are general in the multisensory responding regions. Further, these regions along with primary regions with direct projections or indirectly could also provide a separate only auditory and only visual input source to the OFC.

**Figure 4.**
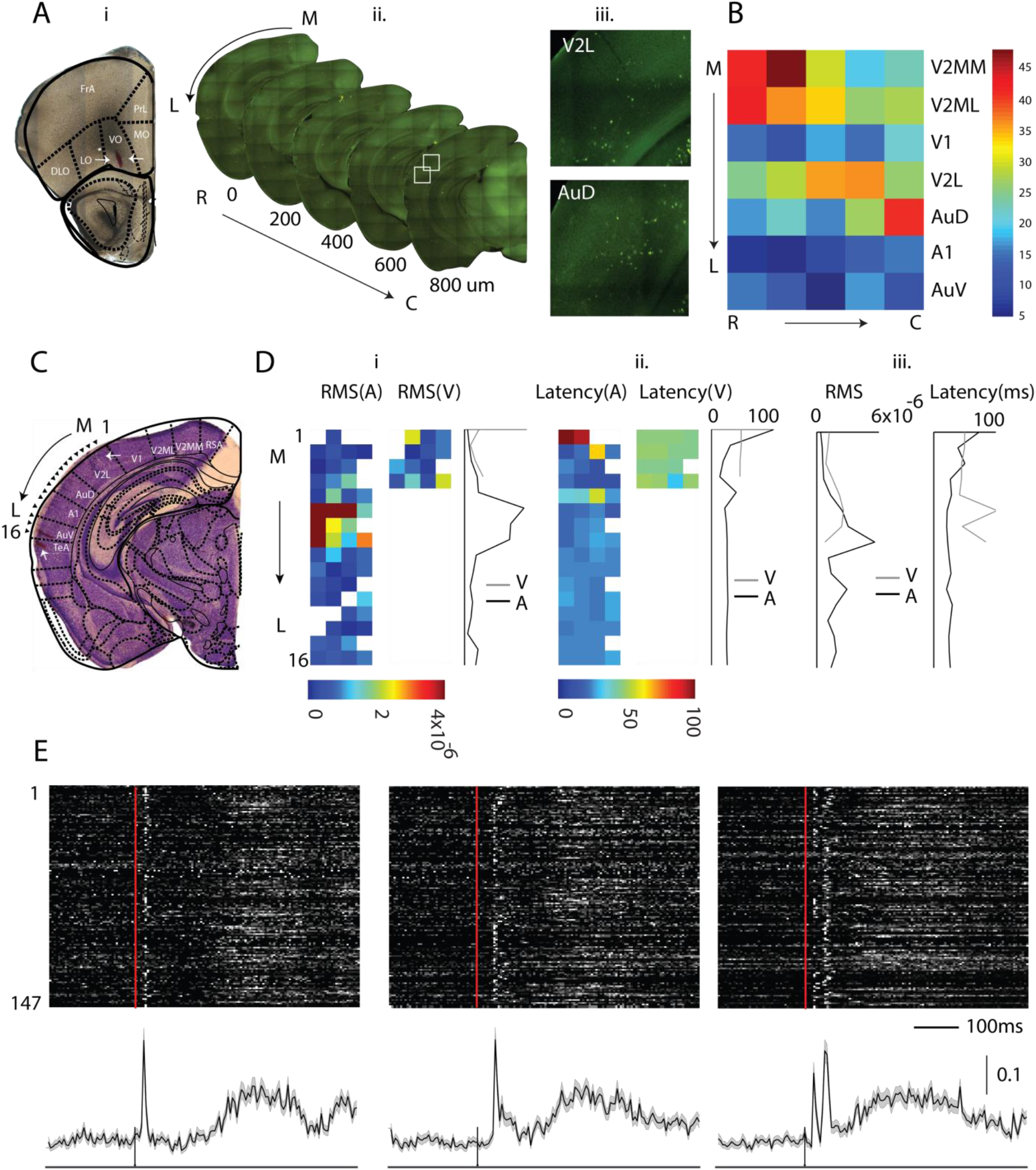
Origin of OFC afferents in the early sensory cortex, carrying auditory, visual and multisensory information. **Ai.** Brightfield image of the frontal cortex section with retro bead injection site (arrow) in OFC. Frontal cortical regions overlaid from mouse atlas(Paxinos and Franklin, n.d.). **Aii.** Example sequence of alternate 100μm fluorescent images of sections encompassing rostrocaudal extent of the ACx. **Aiii.** Sample areas in AuD and V2L (white boxes in Fig 4Aii) with retro bead labeled cell bodies. **B.** Number of labeled cells observed in the different areas. **C.** Nissil stain of an example brain section including the ACx showing electrode tracks (arrows) for recording locations. **Di.** The *rms* amplitude of LFP responses to *A* and *V* (left and middle) for each of the four successive (medial to lateral, ML) recording sites (as in Fig. 4C) with 4×4 MEAs in an example mouse. Right: Mean *rms* amplitude profile along the ML axis for *A* and *V*. **Dii.** Three plots arranged as in Fig. 4Di, for latency of LFP response to *A* and *V* along with the mean profile along the ML axis. **Diii.** Mean ML profiles of *rms* response (left) and latency (right) to *A* and *V* averaged across 4 mice. **E.** Normalized PSTHs of 147 multisensory single units recorded from regions of ACx and VCx immediately dorsal to A1 (*n*=18 mice) shown as rows in grayscale, in response to *A*, V and *M* (left to right). Mean of the normalized PSTHs with *SEM* (gray shading) in each case are shown below. Vertical bar over the population PSTH is the length of normalized rate responses (a.u.).

### Oddball stimulation to parse out modality-specific and multisensory synapses

With clear anatomical connections to the OFC from specific sensory regions, which may underlie the origins of separate *A*, *V* and *M* inputs to the OFC, we designed stimuli to test if that may be the case, from a single unit and LFP responses of OFC. We used an oddball stimulation paradigm with multiple modalities, which makes use of the property of adaptation of synapses with repeated presentations of the same stimulus (called the standard, *S*, Fig. 5A-C), eliminating the response of the postsynaptic neuron (in this case, OFC neurons), while allowing the postsynaptic neuron to still respond to another stimulus (deviant, *D*) that provides inputs through other un-adapted parallel synapses. Complete adaptation to one stimulus thus allows oddball stimulation to look for the presence of different kinds of synapses or input pathways. We used different modality or *M* stimuli in the oddball paradigm to investigate the presence of separate *A*, *V* and *M* channels of input to the OFC single neurons. In all cases (Fig. 5), we consider the normalized PSTHs (normalized by the mean of first 0.5s after stimlus start) of a population of single units. First, repeated presentation of 15 *A*, *V* or *M* tokens at 4 Hz show that neurons in the OFC get completely adapted to each of these stimuli by 1.7s (decay time constants: *A*: 560 ms, *V*: 740 ms and *M*: 80 ms), evident from the rates reaching spontaneous activity levels from the strong onset response. Thus, OFC neurons are completely adapted to the standard stimulus by the end of the 7^th^ token, in such trains of standards (4 Hz).

**Figure 5.**
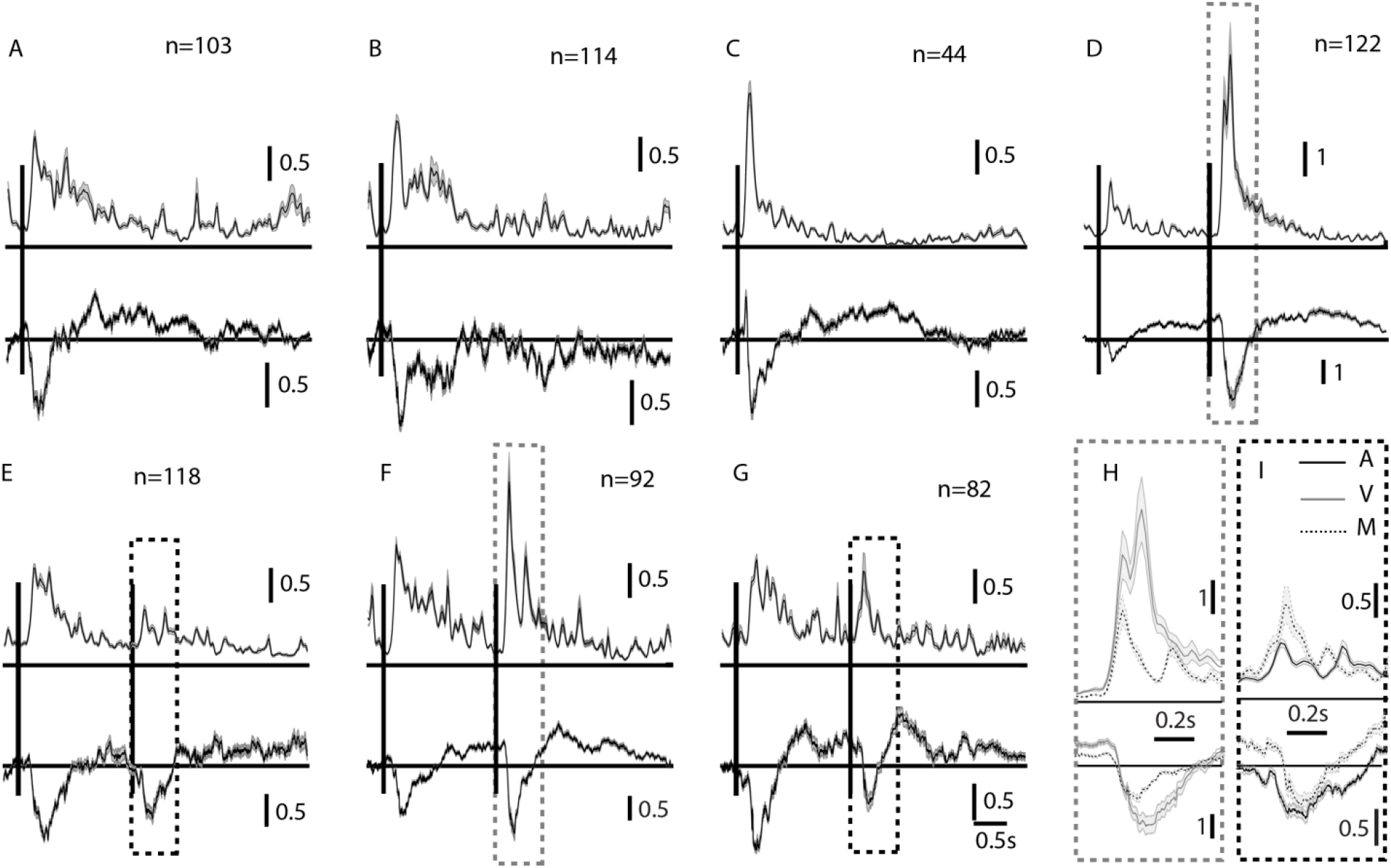
Multisensory oddball stimulation shows the presence of separate *A*, *V* and *M* inputs to the OFC. **A-C.** Mean normalized population PSTHs with *SEM* in response to streams of *A*, *V* and *M* streams of 15 tokens of standards (*S*) Normalized mean LFPs are shown below the PSTHs. **D-I.** Multisensory oddball responses in mean normalized population PSTHs (and LFPs, below) are shown with, *S*=*A* and deviant, *D*=*V*, *S*=*V* and *D*=*A*, *S*=*A* and *D*=*M* and *S*=V and *D*=*M* respectively. Vertical bars show onset (first token) of *S* and the time of *D*. Dashed rectangles (gray, *S*=*A* and black, *S*=*V*) mark the deviant response and pairs of them are compared (in spike rate PSTHs, above and LFPs, below) in H and I respectively.

We use the 8^th^ position, 1.75s from stimulus stream onset, in trains of tokens for the *D* oddball stimulus. In the case of unisensory *S* and *D*, OFC neurons responded to the onset/first of the *S* stimulus tokens (*A* or *V*) and then again to the deviant stimulus (*V*/*A*, Fig. 5D-E). The above results suggest that there are parallel input pathways to OFC that are capable of carrying only *A* and only *V* inputs. The inputs are not necessarily unisensory, but could also be *M* in nature, one responding to *M* and *A* but not *V* and the other responding to *M* and *V* but not *A* or the two inputs could be a combination of unisensory (*V* or *A*) and either of the above (*M*/*A* or *M*/*V* respectively). So to further understand the nature of inputs we used *M* as the *D* token in streams of *A* or *V* as the *S* tokens (Fig. 5F-G). If there are inputs that are of the kind which respond to either one of the unisensory stimuli and *M* then in the case of *M* as *D* in a stream of *A* or *V* there should be no response to the *D*. In both cases the *M* stimulus as *D* evokes a response, indicating that there are exclusively unisensory inputs to the OFC.

Next, we consider if there are separate *M* inputs to OFC, that is inputs that are active only when *M* stimuli are presented and not when unisensory (*A* or *V*) stimuli are presented. We compare the response to *M* as the *D* token, with the response to *V* (or *A*) as the *D* token in a stream of *A* (or *V*) as *S* tokens. This difference in normalized spike rates (7.04±11.86 & 2.69±3.82, *p*<0.001, Fig. 5DF&H and, 1.13±0.70 & 1.85±2.74, *p*<0.01, Fig. 5EG&I) that occur between multisensory deviant and unisensory deviant in the presentation of auditory or visual standards respectively (Fig. 5H&I) indicate the presence of a separate multisensory synapse. Similarly, the comparison in normalized LFPs obtained in each case and their difference (1.49 (IQR=2.39), 0.77 (IQR=0.98) *p*<0.001, Fig. 5DF&H and, 0.45 (IQR=0.98), 0.31 (IQR=0.50), *p*<0.05 Fig. 5EG&I, median compared due to outliers) further indicates that such parallel paths are not emergent in the OFC. In the case of *A* standards, the response to the *V* deviant is larger than the *M* deviant while in case of *V* standards the response to the *M* deviant is stronger than the *A* deviant. This differential effect suggests that there is a multisensory inhibitory input as well to cause the decrease in response to *D*=*M* compared to *D*=*V* in a stream of *A* standards. A comparison of LFPs of the same also suggests that the multisensory inhibition may be driven from outside OFC. Thus as observed based on latencies (Fig. 3A) and with the oddball experiments we conclude that other than separate excitation pathways both *V* and *M* also provide inhibition to the OFC.

### Rapid differential plasticity with audio-visual pairing in the OFC

We find that congruent audiovisual stimulus presentation shows different types of modulatory effects over the single neuron unisensory responses in the OFC. Further, we show that the three stimuli *A*, *V,* and *M* are represented differently with unique response properties based on rates and basic temporal response characteristics. The three stimuli can be decoded or discriminated from each other based on various response temporal features. However, our above results are based on only simultaneous presentations of the two stimuli, while a multisensory stimulus forms its separate identity by continuous passive or active associations. In order to investigate the formation of a multisensory stimulus identity at the circuit level in the OFC, we considered the effect of repeated pairing of *A* and *V* in responses of single neurons to the component *A* and *V* stimuli subsequently, to understand how the separate input pathways deciphered, interact to change representation of the component stimuli.

Following presentations of *A* and *V* stimuli separately (30 repetitions each, Fig. 6A&Bi-ii) to characterize the basic response to unisensory components, 100 repetitions (1 pairing every >7s) of congruent *A* and *V* were presented (Fig. 6A&Biii). Following the pairing responses to *A* and *V* (Fig. 6Biv-v) were again collected to be compared with responses pre-pairing to identify the effects of rapid plasticity, if any. As controls, to rule out the effects of the elapsed time and habituation to the presentations of each of the unisensory components, during pairing, separate experiments were done (Fig. 6A, C-E). For the control experiments, instead of pairings, only the auditory stimulus (*A*=1, *V*=0, Fig. 6D) was presented in one case, only the visual stimulus (*A*=0, *V*=1, Fig. 6E) in another and no stimulus (A=0, V=0) where time equivalent to 100 pairings was given and the spontaneous activity was recorded (Fig. 6C). In each of the control experiments, at the end of the control exposures, responses to *A* and *V* were collected again (Fig. 6C-Eiv&v).

**Figure 6.**
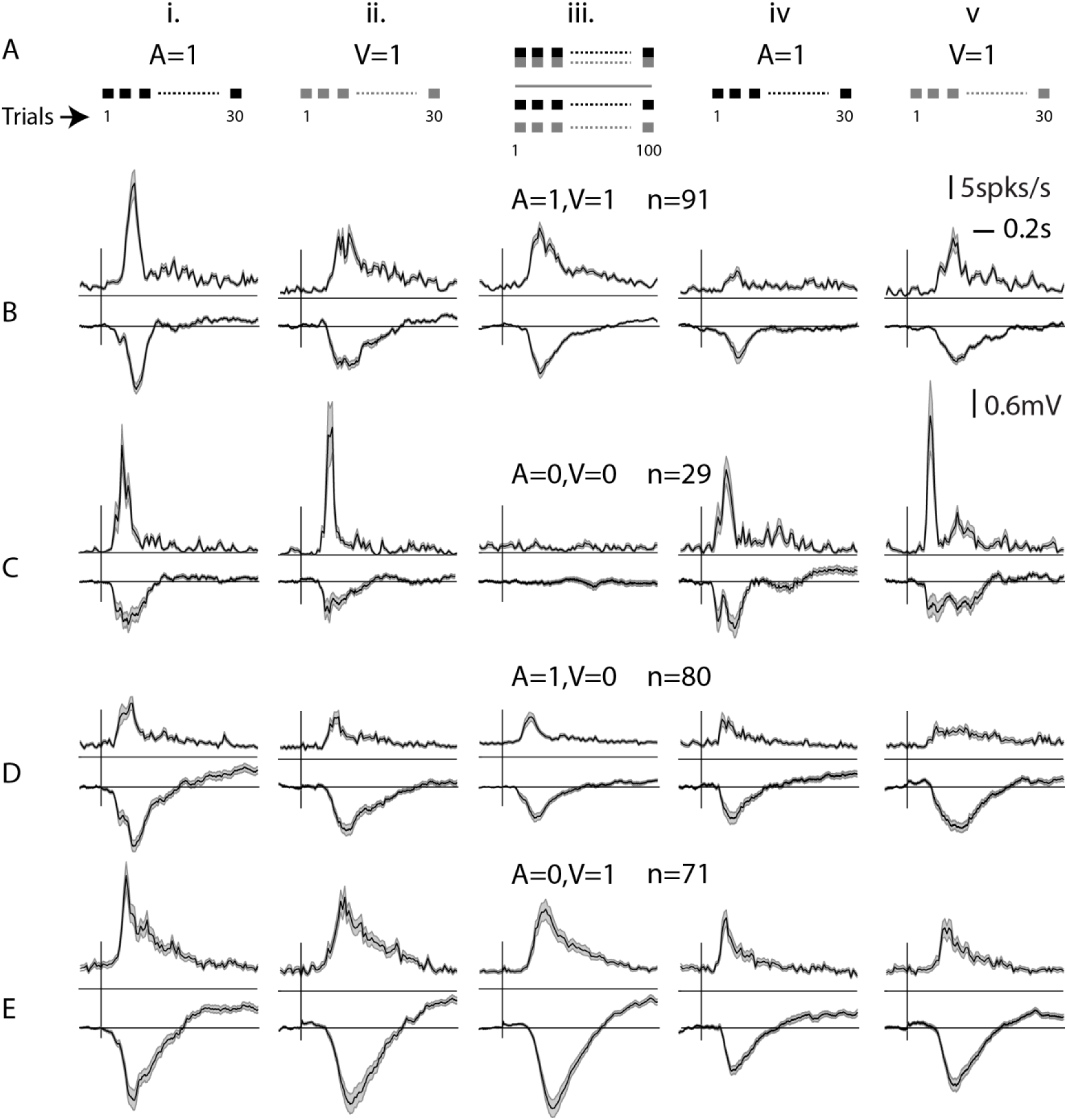
Population effects of audio-visual associations in the OFC. **A.** Experimental protocol is displayed in cartoon form. **B-E.** Each row shows PSTHs with *SEM* of populations of OFC single neurons combined across different experiments. First row corresponds to the case of audiovisual pairing, followed by stability control (no exposure in between), only auditory control and only visual control respectively. First two columns show responses to *A* and *V* before the pairing (or control exposure) and the last two columns show the same after each exposure case. The middle column shows responses during the exposure (100 repetitions or equivalent silence).

In the population of OFC neurons, we observed rapid plastic changes in responses to *A* and *V* due to passive exposure to repeated congruent *A*-*V* stimulation. Following the pairing the spike rate of OFC single neurons in response to both *A* (Fig. 6Biv, Fig. 7Ai, p<0.001) and *V* (Fig. 6Bv, Fig. 7Bi, p<0.01) were suppressed. The suppression of responses to *A* was stronger compared to the mild suppression seen in *V* responses (one-sided comparison of slopes based on bootstrap confidence intervals, Methods, slopes - *A*: 0.1521 and *V*: 0.8071). Significant suppression was also observed in the LFPs following pairing but to different degrees (Fig. 7C-Di, p<0.001 for *A*, slope 0.1885, p<0.001 for *V*, slope 0.5892). Strength of the depression of *A* responses in LFPs was the same as that in spike rates (slopes for before after spike rates 0.1521 and before after LFP *rms* 0.1885, NS, Methods), indicating that the long term depression observed in the OFC auditory response spiking activity is derived from the input pathway. However, there was large variability in the before and after LFP relationship with some LFP channels showing no change or enhancement. The suppression to *V* after pairing was stronger in LFPs than in spike rates (one-sided comparison with bootstraps, Methods) indicating that there is a net enhancement on top of the already suppressed visual activity input to the OFC.

**Figure 7.**
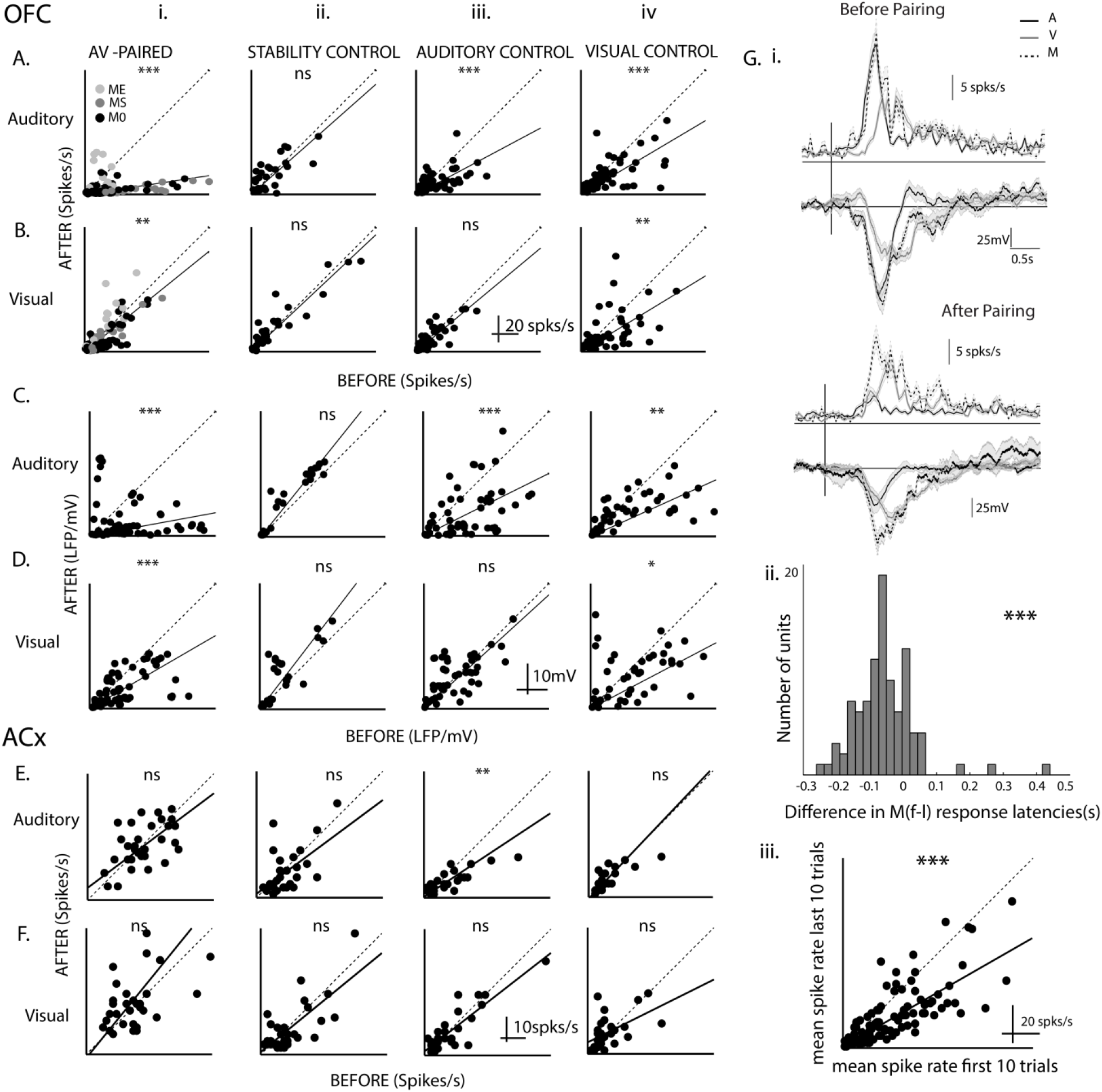
Differential rapid plasticity in *A* and *V* responses with *A-V* association. **A-F.** Each panel is organized in the same way showing scatter plots of after versus before responses of single neurons in spikes (A&B and E&F) or in LFPs at each recording site (C&D) to *A* (AC&E) or *V* (BD&F) stimuli. A-D are for data from the OFC and E&F are for data recorded in AuD/V2L. Each panel has 4 plots (i-iv) starting with *A-V* pairing followed by the 3 control cases. **Gi.** The response to *A* and *V* before (upper) and after (lower) pairing is shown with the population mean PSTHs (above) and mean LFPs (below). Overlaid on them are responses to *M* (before: based on the first 10 trials of pairing and after: based on the last 10 trials of pairing). **Gii-iii.** Distribution of latency differences between first and last 10 trials of pairing response and scatter plot of mean response rate of last 10 trials versus the first 10 trials.

The *ME*, *MS* and *M0* units (Fig. 7Abi, different symbols) showed distinct effects both in auditory and visual responses. The multisensory modulation that was observed, with the audiovisual association changed the responses in a manner so as to make the effect of modulation stronger and longer term. The *ME* units showed relative enhancement of *A* responses compared to the population effect on *A* responses of pairing (*ME* after versus before slope 0.6246 significantly greater than 0.1521, Methods). The effect on *M0* units was not significantly different from the population (*M0* slopes 0.1647) while the *MS* units were significantly more suppressed than the population (*MS* slope 0.1000). Thus if the overall effect of pairing is considered to occur in all units, then the final auditory responses after pairing were determined by the kind of multisensory modulation that operated on the overall effect. A similar effect was seen on visual responses as well following audio-visual pairing. Although *M0* and *MS* units did not show any significant difference in their after versus before relations due reflected in the slopes (*MS* slope 0.7087, *M0* slope 0.6741, NS, Methods), the *ME* units showed a strong enhancement of visual responses (*ME* slope 1.4045 significantly greater than overall slope 0.8071, Methods).

To rule out a general reduction in spike rates over time, control experiments were performed and comparisons were made between responses before and after a time period equivalent to the duration of pairing with no stimulus presentation (*A*=0, *V*=0, Fig. 6C). Neither rate responses to *A* nor that to *V* changed between before and after the no stimulus period (Fig. 7ABii). The same was observed in LFPs (Fig. 7CDii). Thus, the recordings are stable throughout the duration of the experiments and hence the effects seen cannot be due to fluctuations that can occur over time.

To further rule out habituation in auditory responses (or visual responses)(Taaseh et al., 2011) to the repeated presentation of *A* (or *V*), control experiments with 100 presentations of only *A* (*A*=1, *V*=0, or only *V*, *A*=0, *V*=1), instead of pairing, were performed. Habituation based reduction of responses to *A* was observed on repeated *A* presentations (Fig. 6Div, Fig. 7Aiii, p<0.001) and the suppression was weaker than that observed in pairing (before after slopes in case of *A1V1*: 0.1521 and in case of *A1V0*: 0.5370, one-sided comparison with bootstraps, Methods). Thus the repeated presentation of *A* could not explain the long term depression in *A* responses with pairing. Further, the observed suppression in LFPs and that in spike rates in response to *A* after *A*=1, *V*=0 exposure, had no difference, indicating that the habituation observed with only *A* exposure is due to the habituation in the input pathway. Visual responses, both in spike rates and in LFPs, did not change after *A*=1, *V*=0 exposure compared to that before. The *A*=0, *V*=1 exposure produced suppression in LFPs between after and before responses to both *A* and *V*. Similarly, in spike rate responses also, suppression was observed to both *A* and *V*. The suppression in spike rates in response to *V* after *A*=0, *V*=1 is stronger than that after pairing *A*=1, *V*=1 (one-sided comparison of slopes based on bootstrap confidence intervals, Methods, slopes 0.6116 for *A0V1* and 0.8071 for *A1V1*), also further indicating that pairing effectively induces an enhancement in visual responses in the OFC.

Thus from the above experiments, we find that all exposures lead to suppression in the *A* responses, and after pairing, the suppression is stronger than that after only *A* and only *V* exposures, and also stronger than the net multiplicative effect of the individual suppressions (multiplicative slope for *A*, *A1V0xA0V1* 0.3200 higher than *A1V1* slope 0.1521, bootstrap comparison, Methods). On the other hand, the net multiplicative effect on visual responses (multiplicative slope for *V*, *A1V0xA0V1* 1×0.6116) was lower than the slope observed in the case of pairing (*A1V1* slope for *V* 0.8071, bootstrap comparison, Methods). Also, pairing causes suppression in *V* responses in LFPs and hence a net enhancement of the *V* responses over the inputs is observed in the spike rate of OFC neurons. Additionally, with pairing the suppression observed with only visual exposure is overcome. Thus overall we find a differential rapid plasticity of *A* and *V* responses with multisensory pairing. Considering the responses in the last 10 trials of the pairing as the final representation of the *M* stimulus following pairing, we find that the representation of *M* is close to only a visual response showing the visual bias in representation of *M* (Fig. 7Gi). Similarly comparing the latency difference in the first 10 trials and last 10 trials of pairing, shows an increase in latency (Fig. 7Gii, median 60 ms, same as the difference in latency of *M* and *V*, Fig. 3Aiv), also showing the *M* response following pairing to be more like the *V* only response. An effective decrease in overall response is observed (Fig. 7Giii) between the first 10 and last 10 trials of the pairing also consistent with the idea of the *M* response being dominated by the *V* component. The only visual exposure causing the *A* responses to be suppressed (Fig. 7Aiv) with a similar change in LFPs shows that repeated multisensory and visual presentation strengthens the inhibition of the *A* responses by *M* and *V* in the inputs to OFC. Hence the inhibition of *A* by *V* and *M* as hypothesized earlier based on latencies and suppressive multisensory modulation are present in our proposed model in the inputs to OFC itself.

To further understand the effects of pairing, we considered the same experiments to look at effects of the pairing and the related control exposures on *A* and *V* responses in single neurons of AuD/V2L, one of the likely origins of the multisensory inputs (Fig. 4) to the OFC. The same exposure experiments and quantification were performed as done in the OFC (Fig. 7A–8A-D) and the data are summarized in Fig. 8EF). No changes in spike rate responses to *A* or *V* were observed before and after any of the exposures except a suppression in *A* responses after *A*=1, *V*=0 exposure (*p*<0.001). Thus effectively a mild enhancement in *A* responses can be concluded in AuD/V2L with the pairing. Thus the origins of the multisensory inputs to the OFC do not show any of the effects observed in the OFC with pairing, and the changes are derived from higher-order regions. Thus, although similar types and same relative abundance of multisensory modulation was observed in the AuD/V2L and OFC, the effects of pairing to create associations between the *A* and *V* stimuli showed primary effects in higher-order regions than AuD/V2L, including the OFC.

**Figure 8.**
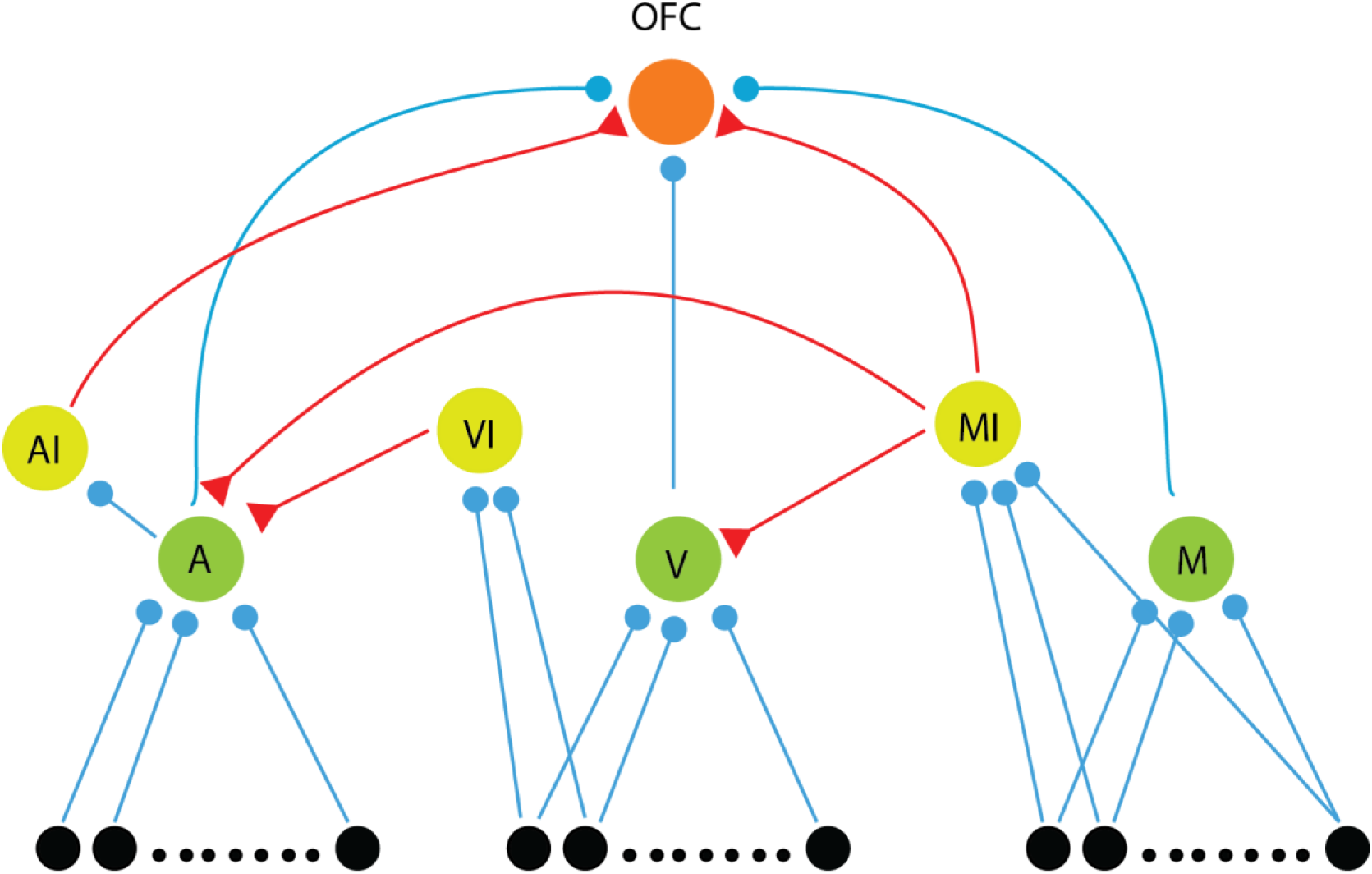
Network model to explain the observations. Hypothetical network model showing inputs to the OFC: 3 excitatory inputs, *A*, *V* and *M* (green neurons, blue synapses) and 2 inhibitory inputs *AI* and *MI* (yellow) neurons with red synapses. *MI* also inhibits the *A* and *V* input neurons and another inhibitory neuron *VI* inhibits *A*. Three separate input populations (black neurons projecting to *A*, *V* and *M*) activated in the presence of *A*, *V* and *M* stimuli respectively (*M* stimulus also activates inputs to *A*and *V*) form the overall inputs to the network.

### Proposed Network Model

Based on our results, we propose a hypothesized minimal network model that qualitatively explains our observations (Fig. 8). We consider the three excitatory input neurons (A, V and M) to the OFC, arising from three separate populations. Three inhibitory neurons are also considered, AI, VI, and MI. The inputs to M are activated by an M stimulus only and not activated with only A stimulus or only V stimulus. The A and V input populations are both activated along with the M input population for the M stimulus. The A neurons providing input to the OFC receive inhibitory inputs from both the inhibitory neurons. The AI neuron provides a weak inhibitory input to the O neuron. The MI neurons inhibit A neurons strongly and the V neurons weakly and also provide inhibitory input to the O neuron.

The MI and VI neurons are lower threshold than V and M neurons to inhibit A with M stimulation. The synapses are considered to be depressive in nature(Mill et al., 2011), which is also the basis for our multisensory oddball paradigm results. With the above, a latency difference as observed can be explained, which allows the O neuron to be responding with the latency determined by the M neuron for M stimulus with A getting inhibited. With the AI neuron inhibiting the O neuron in a delayed manner would allow for the response durations for A to be shorter than that for V and M. Comparatively, in case of the M stimulus, the MI and AI inhibition on O can be overcome by the additional A and V inputs in case of M unlike in case of A, above. Thus M can have longer response duration. With depressing excitatory synapses, the SD stimulus responses can also be explained. Particularly, the decrease in M response compared to V response as deviants in streams of A standards is explained by the MI input to O. For the rapid plasticity observed, with repeated AV pairing, O neurons respond strongly to M inputs inhibiting A, and thus depression in the A->O synapse. As the inhibition from M to V is weak and V still responds during M stimulus, causes enhancement of the V -> O synapse. The above would also lead to the strengthening of the M ->O synapse.

## Discussion

While the OFC has been extensively studied in the context of behavior little is known about the basic response properties of OFC neurons to sensory stimuli, especially in the mouse. In this study, we consider single neuron responses of OFC to auditory and visual stimuli and also to audio-visual (multisensory) stimuli. Our study shows that the multisensory nature of the mouse OFC is in concurrence with other similar studies in the Ventrolateral PFC of primates (Sugihara et al., 2006)(Jung et al., 2018). Comparing responses to component unisensory stimuli and response to the multisensory stimulus(Miller et al., 2015), we find that about equal proportions of neurons to be modulated and otherwise with simultaneous presentation of the unisensory stimuli. The lack of spike rate change in 54% neurons from the maximum unisensory response in case of the multisensory stimulus suggests no multisensory modulation but is, in fact, a nonlinear interaction. Such responses can be explained through extra inhibition based on multisensory stimuli in the OFC (Fig. 8). The other half of neurons showed either enhancement or suppression depending on the strength of responses to the auditory component, showing that auditory inputs guide initial modulation by *M* stimuli. The neurons also were found to encode stimulus identity in specific temporal profiles which are broadly reflected in the latencies and duration of response to different stimuli. The presence of nonlinear interactive effects for the multisensory stimulus even with a 100ms difference between responses to the auditory and visual components shows likely subthreshold mechanisms involved in causing the observed multisensory modulations.

Latency to the three stimuli in this study is in concurrence with previous literature from VLPFC (Romanski and Hwang, 2012). We believe that for audiovisual integration to take place, the generation of a strong prediction in OFC about the late-arriving stimulus elicited by the first-arriving stimulus can cause such latencies to appear (Mazzucato et al., 2017). However, Rowland et al., 2007 suggest multisensory stimulus consisting of simple LED flash and noise results in shortest latencies in the superior colliculus of the cat which could be the result of the subthreshold addition of modality-specific inputs crossing threshold sooner than either of the unisensory stimulation. Hence, the difference may lie in distinct subthreshold properties and receptive fields of the neurons in the two regions.

We show that OFC neurons receive sensory inputs directly from the auditory and visual cortex, more from nonprimary than primary regions, and also possibly indirectly through TeA, Prh, PPC and BLA, which are involved in processing temporal and spatial aspects of visual information and content of auditory stimuli. Thus, OFC has access to both filtered information from auditory and visual specific areas and other aspects of sensory-driven information from higher association areas, representing a redundant multilevel process of integrating information across different sensory systems. Our conclusion of the presence of separate auditory, visual and multisensory inputs based on multisensory oddball experiments supports the above conclusion. Based on the *AV-*oddball stimulation results, we find there are effective parallel unisensory (*A* and *V*) and multisensory (*AV*) inputs to the OFC and the findings are summarized with additional components to explain our other results, in the proposed network model (Fig. 8). It is to be noted that we consider effective pathways, in the sense that there are likely multiple parallel *M* inputs and similarly *A* and *V* inputs. A further detailed network model would involve parsing out each one of them through pharmacological or optogenetic manipulations (Deisseroth et al., 2006). While multisensory audio-visual modulations are known to play a role in primary as well as secondary auditory and visual cortices (Hollensteiner et al., 2015)(Bizley and King, 2008)(Meredith et al., 2009) in sensory coding, as in our AuD/V2L responses, we find that similar modulations in similar proportions of neurons to also exist in the OFC. Thus such multisensory modulations are likely intrinsic in the sensory pathways.

However, in terms of multisensory interactions, the most notable difference in multisensory processing observed between sensory cortices and the OFC is the outcome of multisensory associations or audio-visual pairing. We determined the effect of creating multisensory associations and their subsequent effect on unisensory stimulus processing in the OFC and AuD/V2L. We found differential plasticity in auditory and visual responses on long passive associations between the auditory and visual stimulus in the OFC. Such association driven plasticity was absent in the AuD/V2L single neurons. A visual bias was observed after multisensory associations with strong suppression of *A* responses and effective enhancement of *V* responses. Such differential rapid plasticity is intrinsic to the circuitry in OFC and independent of reward, behavioral context or other cognitive aspects. The above result shows that the OFC circuitry intrinsically has a bias towards weighing visual stimuli differently from auditory stimuli in such associations when in behaving conditions the OFC would engage in multisensory tasks. Similar biases to olfactory or particular features of sensory stimuli have been observed (Kuang and Zhang, 2014), however, such sensory bias in associations absent in the sensory cortex and found in the higher cortex is not known. Further, since the OFC is also capable of modulating early sensory responses (Winkowski et al., 2013), such differential plasticity in the OFC can thus be critical in determining stimulus representation in the early sensory cortices in a multisensory environment.

Since the responses in the OFC are heavily dependent on behavioral state, memory, and context, in order to understand the intrinsic response properties to sensory stimuli in the OFC, we have considered the responses in an anesthetized preparation. While anesthesia, in this case isoflurane, is known to alter responses through increased GABAA receptor activation and NMDA receptor inhibition (Capey, 2007). In the OFC, we consider that the basic observations in our study would remain intact in terms of the properties of the circuits leading to sensory responses. Further, the results of rapid plasticity would also likely remain unchanged as NMDA receptor inhibition with isoflurane suggests stronger plastic changes to be expected without anesthesia.

Oddball stimulation has been used previously to study different frequency channels for probing frequency encoding in the auditory cortex (Taaseh et al., 2011) and the effects can be explained by parallel channels of depressive synapses (Mill et al., 2011). In our study, we adapted it to study input channels from different sensory modalities, which in our case, very clearly reveals the presence of separate auditory, visual and multisensory inputs into OFC. Based on the results of the oddball stimulation, we could also conclude the presence of an inhibitory input that is driven by multisensory stimuli to the OFC. It is to be further noted that recurrent connections, absent in our proposed hypothetical model (Fig. 8), can potentially generate and enhance the effects we have observed (Yarden and Nelken, 2017) in our oddball experiments. We have considered LFPs in parallel with spiking responses to loosely provide information on summed synaptic input activity and output activity respectively. Such comparisons as indicated also corroborate our conclusion of parallel input pathways for the 3 types of stimuli. If there are subthreshold depolarizations caused by inputs from synapses which we assume to be completely adapted with a sequence of standard presentations, then our conclusions need reconsideration. LFPs, known to indicate summed synaptic activity, however, do not show any evidence of synaptic inputs on presentations of repeated standards beyond the first two to three tokens.

Our finding of the OFC in the mouse being primarily multisensory (audio-visual) was not known before. The sensory inputs to the mouse OFC mostly studied have been unisensory in nature and primarily the chemical senses associated with food. Although responses in the mouse OFC to auditory stimuli (Winkowski et al., 2017) are known, there have been almost no previous studies of mouse OFC responses to visual stimuli or audio-visual stimuli. Thus, our findings are critically important for understanding how sensory responses are dynamically modified to achieve a stable perception of a dynamic environment and how changing value based on the behavioral relevance of incoming stimuli may be computed. Most importantly our findings indicate an intrinsic bias towards visual components over auditory components in the representation of multisensory stimuli in the OFC. Such a differential effect on one sense over the other would be critical in multisensory association based behavior and value computation and similarly in adjusting sensory representations in a dynamic multisensory environment. Since the OFC is capable of modifying sensory responses (Winkowski et al., 2013) such a visual bias, can cause behaviorally relevant visual components of multisensory stimuli to modify auditory responses during reward-based learning.

